# Brain Connectivity meets Reservoir Computing

**DOI:** 10.1101/2021.01.22.427750

**Authors:** Fabrizio Damicelli, Claus C. Hilgetag, Alexandros Goulas

## Abstract

The connectivity of Artificial Neural Networks (ANNs) is different from the one observed in Biological Neural Networks (BNNs). Can the wiring of actual brains help improve ANNs architectures? Can we learn from ANNs about what network features support computation in the brain when solving a task?

ANNs’ architectures are carefully engineered and have crucial importance in many recent performance improvements. On the other hand, BNNs’ exhibit complex emergent connectivity patterns. At the individual level, BNNs connectivity results from brain development and plasticity processes, while at the species level, adaptive reconfigurations during evolution also play a major role shaping connectivity.

Ubiquitous features of brain connectivity have been identified in recent years, but their role in the brain’s ability to perform concrete computations remains poorly understood. Computational neuroscience studies reveal the influence of specific brain connectivity features only on abstract dynamical properties, although the implications of real brain networks topologies on machine learning or cognitive tasks have been barely explored.

Here we present a cross-species study with a hybrid approach integrating real brain connectomes and Bio-Echo State Networks, which we use to solve concrete memory tasks, allowing us to probe the potential computational implications of real brain connectivity patterns on task solving.

We find results consistent across species and tasks, showing that biologically inspired networks perform as well as classical echo state networks, provided a minimum level of randomness and diversity of connections is allowed. We also present a framework, *bio2art*, to map and scale up real connectomes that can be integrated into recurrent ANNs. This approach also allows us to show the crucial importance of the diversity of interareal connectivity patterns, stressing the importance of stochastic processes determining neural networks connectivity in general.

## Introduction

Recent breakthroughs in Artificial Neural Networks (ANNs) have prompted a renewed interest in the intersection between ANNs and Biological Neural Networks (BNNs). This interest follows two research avenues: improving the performance and explainability of ANNs and understanding how real brains compute [1].

Many recent improvements of ANNs rely on novel network architectures, which play a fundamental role in task performance [2, 3]. In other words, such connectivity patterns allow for better representation of the outer world (i.e., the data) and/or they let the networks learn better, e. g., promoting faster convergence. Nevertheless, ANNs employ architectures that are not grounded in empirical insights from real brains network topology. For example, ANNs do not follow ubiquitous organization principles of BNNs, such as their modular structure, and BNNs are also much sparser than ANNs [1,4].

Given that Biological Neural Networks (BNNs) present complex, non-random connectivity patterns, it is hypothesized that this “built-in” structure could be one key factor supporting their computation capabilities. In consequence, a focus on BNNs’ topology has started to gain traction in recent ANNs research [5,6]. For instance, building feedforward networks based on graph generative models, such as Watts-Strogatz and Barabási-Albert models, has resulted in competitive performances compared to optimized state-of-the-art architectures [7]. In a complementary vein, feedforward networks may spontaneously form non-random topologies during training, such as modular structure [8]. In addition to that, combining evolutionary algorithms with artificial neural networks has shown that a modular topology can improve performance and avoid forgetting when learning new tasks [9]. In sum, current evidence supports the notion that non-random topologies can lead to desired performance ANNs.

However, studies thus far have only focused on network topology models that have almost no direct correspondences (or only abstract ones) to BNNs unravelled by experimental connectomics. Hence, it is to date unknown if and to what extent the *actual, empirically discerned* topology of BNNs can lead to beneficial properties of ANNs, such as more efficient training (fewer epochs and/or samples) or better performance (e.g., higher test accuracy).

A complementary view comes from *connectomics* and *network neuroscience*, fueled by experimental advances for mapping brain connectivity to an unprecedented level of detail [10]. In that context, a *connectome* refers to all mapped connections of one individual brain, either coming from one individual or aggregated across sampled brains, depending on the experimental methodology. Graph-theoretical tools are then leveraged to describe brain connectivity and find potential associations. For example, looking for correlations between specific graph properties and cognitive tasks performance [11]. Along those lines, some graph properties typical of real brains can also have advantageous dynamical properties, such as supporting the balance between information segregation and integration [12,13]. Nevertheless, the relationship (if any at all) between those abstract dynamical properties and the performance of the network on concrete tasks remains unclear.

We explicitly address that gap here by building recurrent Echo State Networks (ESN) that are *bio-instantiated*, thus *BioESNs*. We ask if and to what extent the topology of BioESNs affects its performance on concrete memory tasks. We build BioESNs that embody the wiring diagram empirically found in brains of three primates species, including humans. We also present a framework, the *bio2art* [14], to map and scale up real connectomes, allowing to integrate them into recurrent ANNs.

This is a necessary step exploring the possible links between biological and artificial neural systems, not by means of abstract network models but exploiting the wealth of empirical data being generated, which has started to paint a detailed picture of the intricate wiring of biological neural networks.

## Results

In order to test the potential effect of the topology of real connectomes on the computation capacities of recurrent networks, we devised a hybrid Echo State Network (ESN) integrating real brain connectivity, thus *BioESN*. Classical ESNs have an internal reservoir of neurons sparsely and randomly connected. The reservoir works as an internal non-linear projection of the input(s), generating a rich variety of features, such that the readout layer can linearly separate the patterns more easily. Thus, the performance of an ESN is related to the richness of the representation generated by the reservoir neurons, in turn related to the connectivity pattern between the reservoir neurons.

We investigate here how the non-random topology of biological neural networks affects the performance of ESN by integrating real connectomes as reservoir and letting the BioESN solve concrete cognitive tasks (see Fig. 1). Importantly, we construct the BioESN reservoirs based on the wiring diagrams (i.e., who connects to whom) of connectomes, but using the weights initialization typically used in classical ESNs (see Fig. 2 and Methods for details).

**Fig 1.**
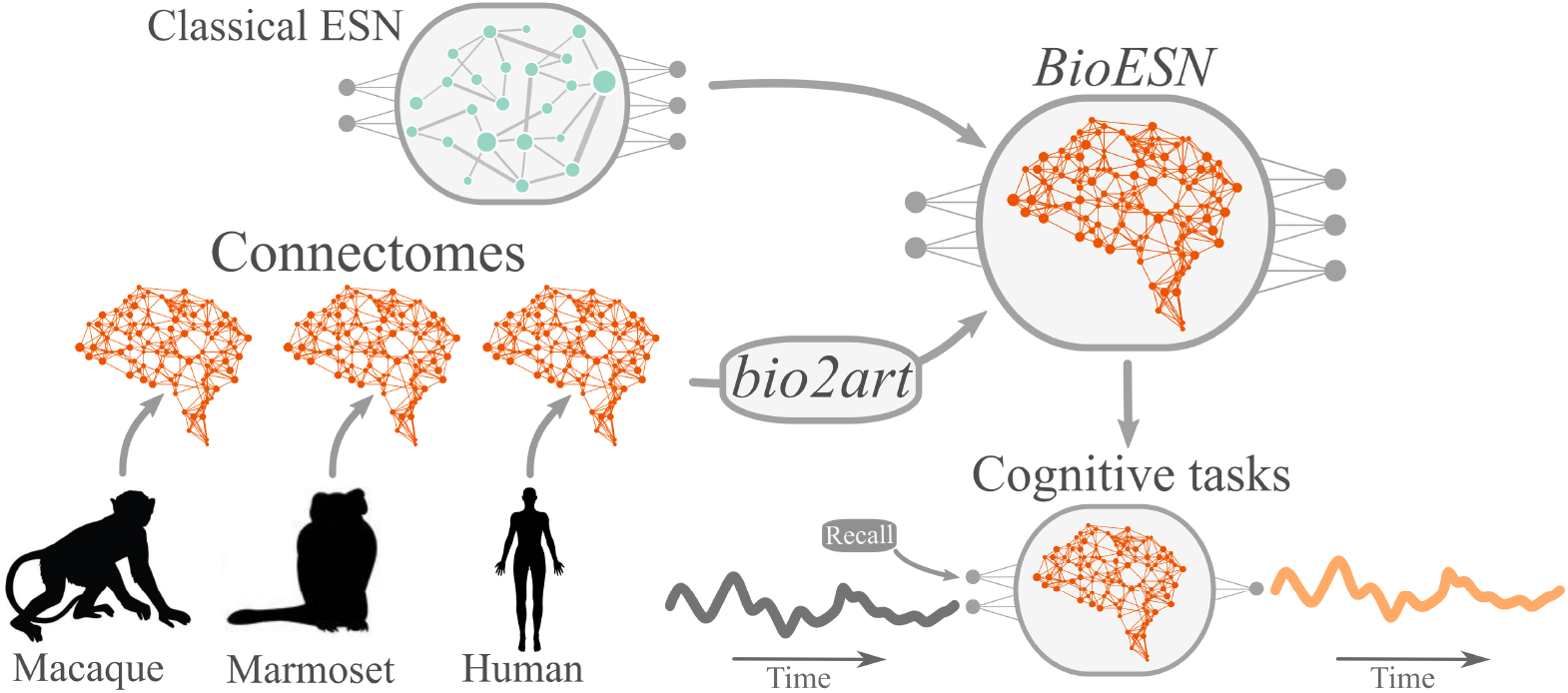
General approach scheme. For each of the three species we generated a Bio-Echo State Network (*BioESN*) by integrating the real connectivity pattern as reservoir of an Echo State Network (ESN). Thus, in contrast to the classical ESN with randomly connected reservoir, BioESNs have connectomes based on connectivity coming from the empirical connectomes. We also propose a framework for mapping biological to artificial networks, *bio2art*, which allows to optionally scale up the empirical connectomes to augment the model capacity. The resulting BioESNs are then tested on cognitive tasks (see Fig 6 and Methods for details on the tasks).

**Fig 2.**
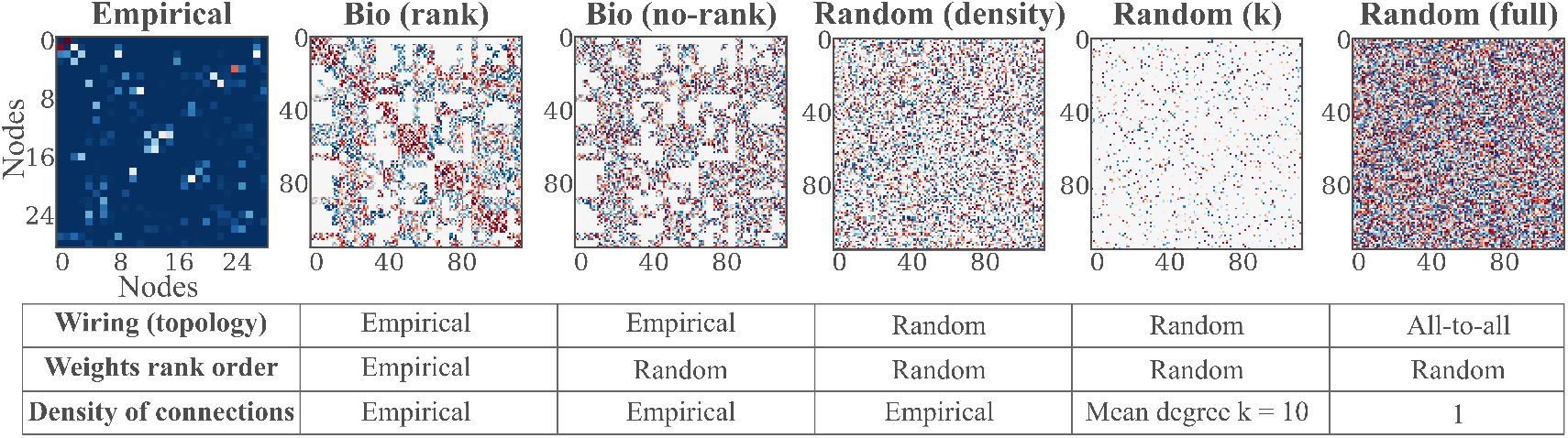
*bio2art*, scaling up connectivity and surrogates. The connectivity of the networks derived from the empirical connectivity and used as reservoirs in the BioESNs can be represented as an adjacency matrix. This figure shows examples of adjacency matrices representing a scaled up version (4x) of the Macaque monkey empirical brain connectivity then integrated into the BioESN as reservoir. We also build surrogate connectivities for comparison with the empirical case that preserves real connectivity patterns. Each surrogate network controls for different aspects of the connectivity, as shown in the summary table in the figure. The figure depicts an example of the empirical (Macaque) connectivity and the different derived connectivities tested. Notice the nodes indices, explicitly showing the upscaling of the connectivity. This was repeated for all the other connectomes tested. See Mappping and upscaling connectomes with *bio2art* for more details on connectivity generation and surrogates.

We tested connectomes of three different primate species (Human, Macaque and Marmoset) in two different memory tasks.

Since we aimed at testing if the connectivity pattern of empirical connectomes could have an effect on the performance of BioESNs, we generated several variations of the connectivity as a surrogate network for contrast, where each surrogate preserves (or not) specific connectivity properties, as summarized in Fig 2.

The conditions *Bio (rank*) and *Bio (no-rank*) preserved the empirical binary topology mask (i.e., who connects to whom) and thus constitute the conditions that we mainly aimed at testing. The difference between both conditions is that *Bio (rank*) preserved the ranking of the weights. That means, in spite of the weights coming from a random distribution, the links in the network were allocated such that links with high strength in the empirical connectome also corresponded to stronger weights in the BioESN. In contrast to that, such rearrangement of links was not performed for the *Bio (no-rank*) condition, thus only keeping the binary mask of empirical connectivity but respecting no ranking order of links. The other surrogate conditions (*Random (density), Random (k), Random (full*)) have totally random wiring diagrams and their density of connections is the only factor varying across conditions. For the condition *Random (density*), the density of connections is the same as for the empirical connectome. For the condition *Random (k*), the network forming the reservoir has a fixed number of links per node *k* = 10, as in classical ESNs approaches [15]. For the condition *Random (full*), there are no restrictions in terms of links, thus generating a fully connected network, i.e., density equal to 1 (again, please refer to Fig 2 for a summary on all the tested conditions). Larger reservoirs are *per se* expected to have better performance than otherwise equivalent networks [15]. Thus, given that the different sizes of the empirical connectomes, the results are not comparable across connectomes, but only across connectivity conditions for one connectome.

### Memory Capacity Task

In this classical paradigm, the network is presented with a random sequence of numbers through a unique input neuron. Each output neuron is trained independently to learn a lagged version of the input, thus there are as many output neurons as lags to be tested [16]. The performance, the so called *Memory Capacity (MC*), is calculated as the cumulative score across all outputs (i.e., all time lags, see Fig 6 for details).

**Fig 6.**
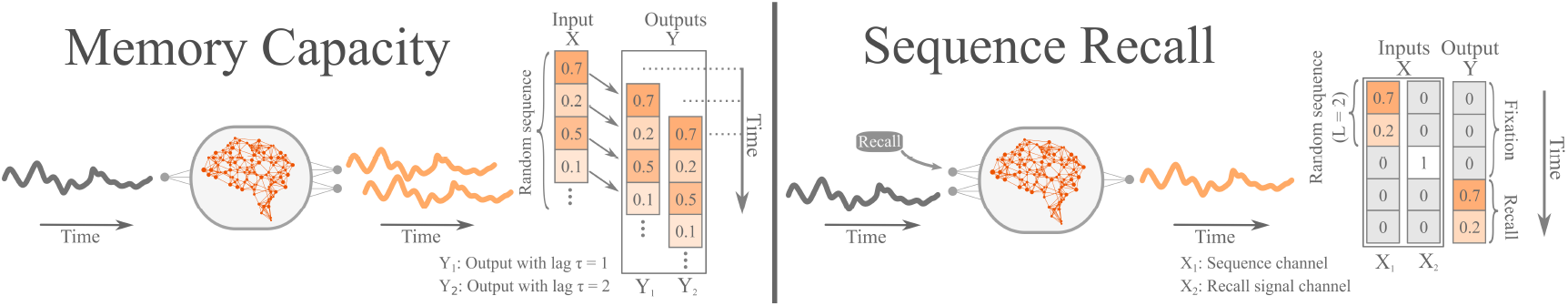
Cognitive tasks. Schematic representation of the tasks and the input/output data structure for each of the cognitive tasks used to evaluate the performance of the BioESNs. Left: Memory capacity (MC) task, where the network receives a stream of random values as single input X and has several independent outputs Y (for simplicity, the example shows only two. Each output is memorized by an independent output neuron of the network and is supposed to recall the input at a specific time lag *τ*. The BioESNs were trained with 4000 time steps and tested on the subsequent 1000. Right: One trial of the sequence recall task. The network receives inputs *X*_1_, *X*_2_ coming from a random sequence and a recall signal channel, respectively. There is only one output neuron, which after the recall signal channel indicates it (i.e., X2 = 1) is supposed to reproduce the input received in the previous L steps, i.e., the pattern length parameter determining the difficulty of the task (for simplicity, in the scheme L = 2). The BioESNs were trained with 800 trials and tested on 200 trials. The score was computed considering only the recall phase in order to avoid inflation of the metric, given that the fixation periods were much easier to perform correctly.

Our results show a comparable performance across all reservoir types except for the *Bio (rank*) condition (Fig 3). Networks from all the tested conditions were able to learn the task, at least for the lowest difficulty (time lag *τ* = 5), but the *Bio (rank*) condition showed significantly worse memory capacity. This pattern was consistent for all the empirical connectomes tested (Macaque, Marmoset and Human).

**Fig 3.**
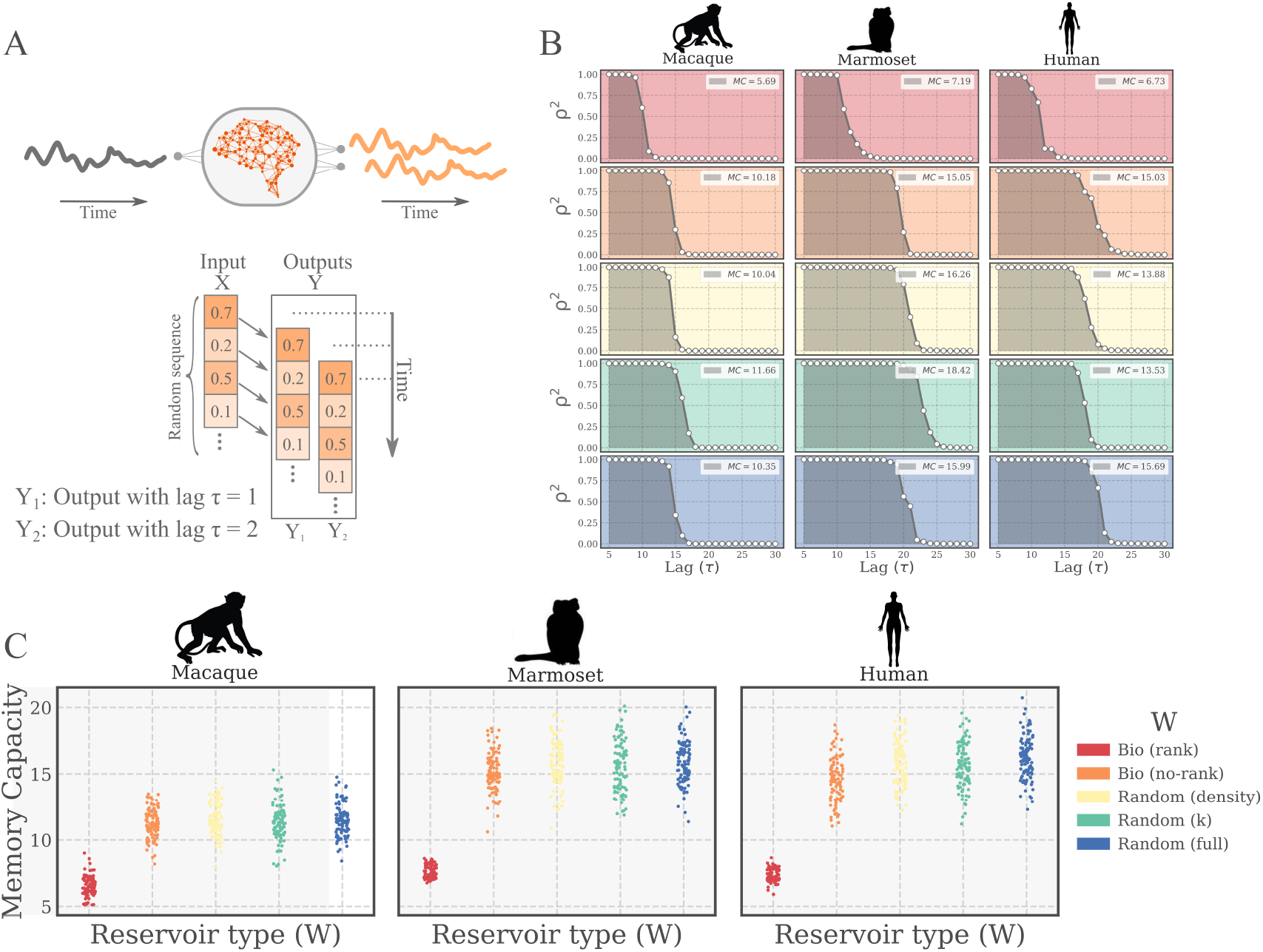
Memory Capacity Task. **(A)** (Upper) Schematic representation of the task. An input signal (*X*) is feed as a time series into the network through an input neuron. Each output neuron independently learns a lagged version of the input (*Y_τ_*) (Lower) Alternative representation of the task in terms of the input/output structure of the data. **(B)** Examples of network evaluation on the task. A forgetting curve (grey line) is shown for each tested species (columns) and connectivity *W* condition (color coded). For each time lag (*τ*) the score is plotted (squared Pearson correlation coefficient, *ρ^2^*). The memory capacity (MC, see legends) is defined as the sum of performances over all values of *τ* and represents the shaded areas in the plotted examples. **(C)** Performance of the bio-instantiated echo state networks (*BioESNs*) for the three different species tested. For each pattern length, 100 different networks with newly instantiated weights were trained (4000 time steps) and tested (1000 time steps). The test performance of each networks is represented by a point in the plots.

On the other hand, we did not find differences in the performance for all the other tested conditions. This indicates that a certain level of randomization is actually necessary to reach a better performance and that biological wiring diagrams can achieve the same performance as the purely random networks, provided an adequate level of randomness is allowed, as in the *Bio (no-rank*) condition.

### Sequence Memory Task

In this task, the network is presented with two inputs, a sequence of random numbers to memorize and a cue input, thus having two input neurons. The cue input indicates whether to fixate (output equal to zero) or to recall the presented pattern. When the recall cue is presented, the network is supposed to output the memorized sequence in the previous *L* steps, where *L* is the *pattern length*, a parameter regulating the task difficulty (see Fig. 6 for details). One trial of the task consists of a fixation period followed by a recall period. In order to avoid inflation of the score, the performance was evaluated exclusively during the recall steps, which are more difficult to perform than the fixation phase.

In agreement with the results for the Memory Capacity task, we found that all tested ESNs were able to learn the task, at least in its easier variations (pattern length L = 5). Along the same lines, the *Bio (rank*) condition was the only one with a significantly different performance, showing worse performance than the rest of the conditions. The *Bio (rank*) networks had a different performance decay profile, only being able to memorize shorter sequences than the rest of the network types (Fig. 4). These findings were again consistent across all the different connetomes tested.

**Fig 4.**
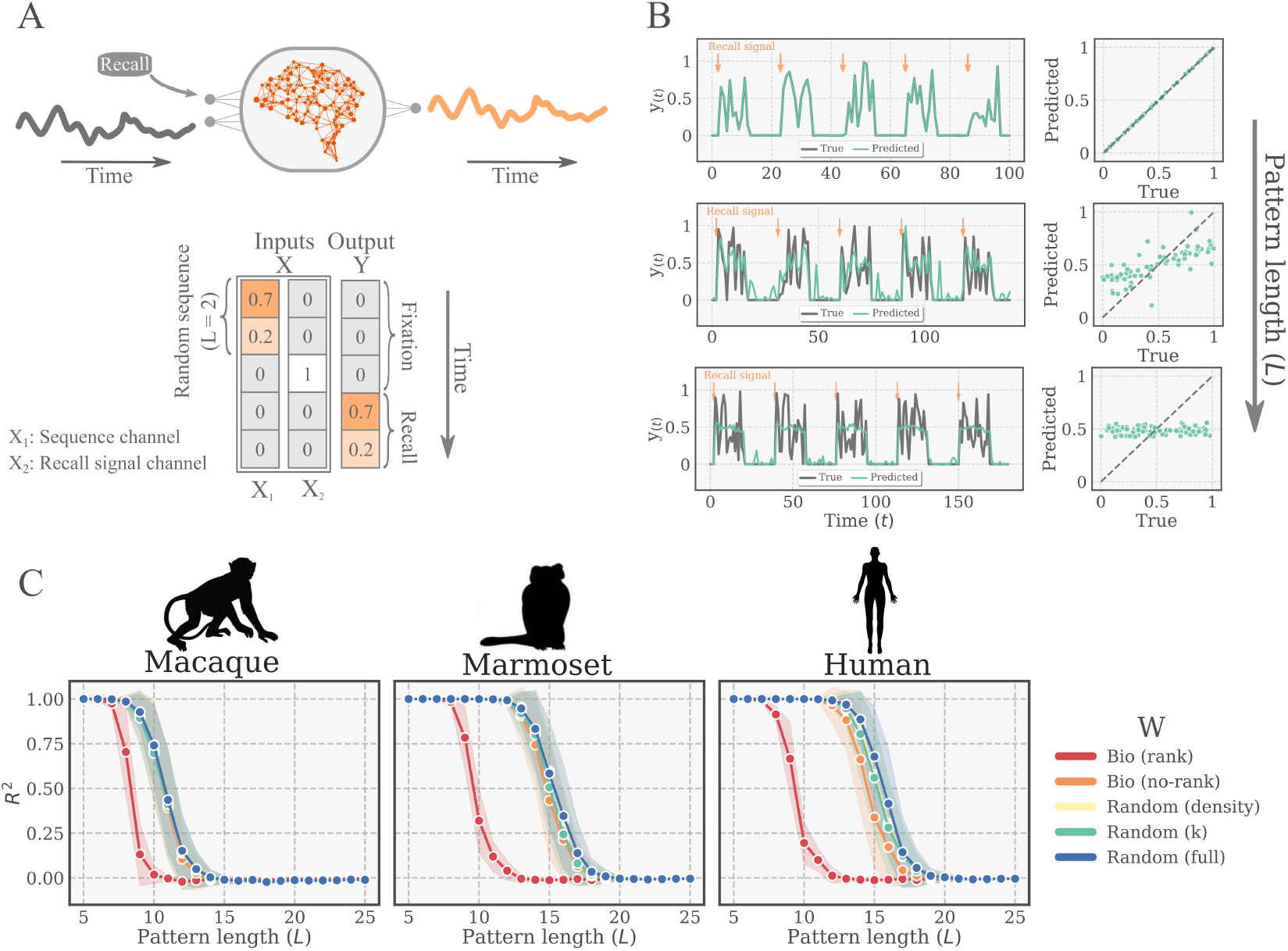
Sequence Memory Task. **(A)** (Upper) Schematic representation of one trial of the task. The input signal (*X*_1_) and the recall signal (*X*_2_) are feed as a time series into the network through two input neurons. When the recall signal is given (*X*_2_ = 1), the output neuron is supposed to deliver the memorized input of the last *L* steps. (Lower) Alternative representation of the task in terms of the input/output structure of the data. **(B)** Examples of actual and predicted times series for 5 trials at three different difficulty levels (pattern length, from top to bottom: L = 10/14/18). The scatter plots on the right show the predicted *vs*. the true output (as explained in main text). The BioESN in the example was built from human connectome with the *Bio (no-rank*) variation. **(C)** Performance of the bio-instantiated echo state networks (BioESNs) for different task difficulties (pattern length) for the three different species. The bio-instantiated reservoirs, *Bio (rank/no-rank*), are compared to surrogates with random connectivity patterns. For each pattern length, 100 different networks with newly instantiated weights were trained (800 trials) and tested (200 trials). The curves depict the mean test performance and standard deviation across networks.

#### *bio2art*: Mapping and up-scaling connectomes

The performance of an echo state network is intimately related to the reservoir size, since a larger reservoir can potentially generate a richer repertoire of features and has more trainable parameters, provided a proper weights initialization. Our next goal was to investigate the scaling behaviour of our model, i.e., how the performance changes as the reservoir size is increased. For that, we applied our *bio2art* approach [14], which allows us to map the connectivity of real connectomes onto artificial recurrent networks and scale up the number of neurons by an arbitrary scaling factor while preserving the wiring diagram of the original connectome (see Mappping and upscaling connectomes with *bio2art* for more details). Although not totally conclusive because of the different experimental methodologies, that approach brings us closer to a comparison across species connectomes (see Discussion). At the same time, this allows us to explore the extent to which the pattern observed for the different weights mappings (Bio (*rank), Bio (no-rank*), etc.) holds for larger networks and how it plays out with the model capacity driven by reservoir size.

When upscaling the connectomes with *bio2art*, one important parameter is whether to generate a homogeneous or heterogeneous distribution of weights between scaled-up areas, as shown in Fig. 5A. With the homogeneous variation, all connections between two upscaled areas have exactly the same weight. In other words, after scaling up the number of neurons per area, the total original weight between every two areas is equally partitioned amongst all the area-to-area connections. In contrast to that, the heterogeneous variation allows the area-to-area connections between scaled up regions to be different. More specifically, the total original weight between every two upscaled areas is partitioned and distributed at random amongst the area-to-area connections.

**Fig 5.**
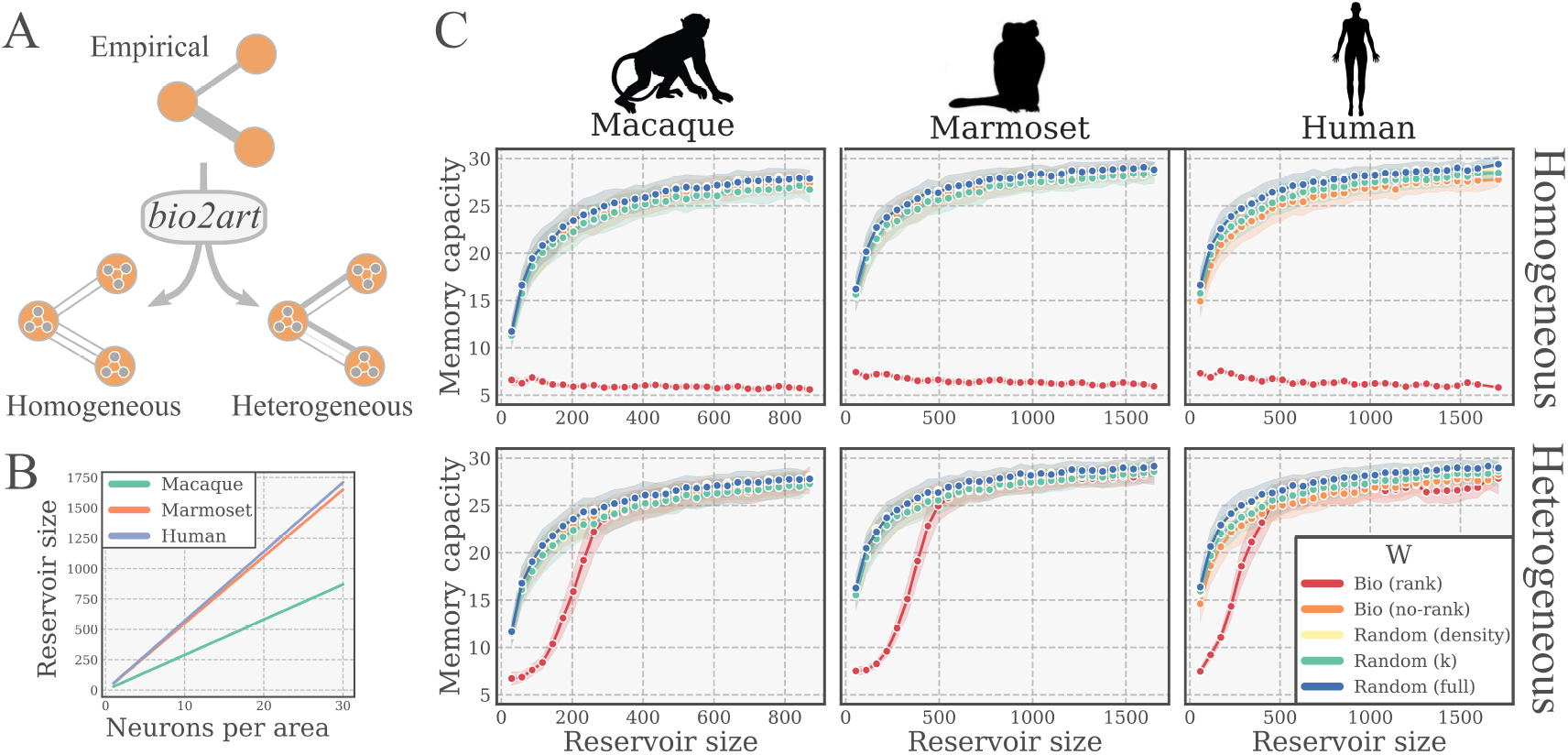
Scaling of performance with reservoir size. **(A)** Scaling up empirical connectomes with *bio2art*. Scaling allows to specify a number of neurons per area (brain region as defined in the connectome). The interareal weights might be mapped either homogeneously or heterogeneously. Homogeneous mapping partitions total weights in equal parts amongst interareal connections. Heterogeneous mapping partitions total weights at random amongst interareal connections. **(B)** Relationship between the neurons per area and the total reservoir size for all the studied scaling factors. **(C)** Performance of BioESNs with scaled up connectomes on the memory capacity task, for heterogeneous and homogeneous interareal connectivity patterns (upper and lower row, respectively). For each single condition (size, interareal connectivity), 100 different networks with newly instantiated weights were trained (4000 time steps) and tested (1000 time steps). The curves depict the test performance mean and standard deviation across runs.

We evaluated a wide range of scaling factors (i.e., neurons per area) for the BioESNs generation with *bio2art*, considering both homogeneous and heterogeneous interareal connectivity pattern as explained above. We found a clear pattern of improving performance, reaching an asympthotic value roughly comparable across all connectomes when looking at reservoirs of same size. Interestingly, the otherwise consistently lower performance of the *Bio (rank*) condition could be reverted with large enough reservoirs. Importantly, this was only the case for scaled up connectomes with heterogeneous interareal connectivity patterns but not with homogeneous patterns (see Fig. 5). This indicates that the randomness and diversity of connections in interareal connectivity plays a crucial role determining the memory capacity of the network. For this series of experiments we also found consistent results across all studied connectomes.

## Discussion

We address two fundamental questions aiming at bridging the gap between artificial and biological neural networks: Can actual brain connectivity guide the design of better ANNs architectures? Can we better understand what network features support the performance of brains in specific tasks by experimenting with ANNs? Concretely, we investigate the potential effect of connectivity built based on real connectomes on the performance of artificial neural networks. To the best of our knowledge, this is the first cross-species study of this kind, comparing results from empirical connectomes of three primate species.

The gap that we aim at emerges from two under-explored aspects in artificial and biological neural networks. First, connectivity patterns (i.e., architectures) of ANNs are very different from actual brain connectivity. For example, echo state networks use a sparse, randomly connected reservoir, which is incongruent with the highly non-random connectivity empirically found in the brain [4,11]. Thus it is not clear, how more realistic architectures would impact the performance such ANNs. Second, computational neuroscience studies have characterized the relation between structural and function connectivity patterns [17, 18] and attempted to relate brain connectivity to behavioural differences [19,20]. Nevertheless, it remains unclear how those patterns of neural activity translate into brain computational capabilities, i.e., how they support performance of brain networks on concrete tasks. We set out to evaluate real whole brain connectomes on specific tasks, in order to identify a potential role of such wiring patterns, in a similar vein to previous studies on feedforward networks [21].

We found that constraining reservoir connectivity of ESNs with real connectomes led to performances as good as for the random conditions, classically used for ESNs, as long as a certain degree of randomness is allowed.

In general, we observe a degeneracy of structure and function, in which different topologies lead to the same performance, so no unique connectivity pattern appears necessary to support optimal performance in this modeling context.

Our results were similar across tasks. This is to a certain extent logical considering that both tested tasks are memory tasks, but the consistency also speaks for the robustness of the networks to different recall mechanisms.

Importantly, all our results were consistent across the three evaluated species. This supports the generality of our findings, at least for the evaluated tasks. This observation is especially relevant considering that the connectomes were obtained with very different experimental methodologies [10,22,23]. Moreover, our experiments with scaled up connectomes showed similar performance scores across species when the reservoir size was matched. Nevertheless, the different experimental methodologies to infer the connectivity prevent us from drawing specific comparative conclusions across connectomes, such as whether the wiring diagram of any of the tested connectomes is intrinsically better suited for the task regardless of the size.

Our surrogate networks also showed that, in general terms, the more heterogeneity and randomness allowed in the connectivity, the better performance the BioESNs achieved. Interestingly, that effect was also observable by augmenting the computational capacity of the models by means of larger reservoirs. Using the *bio2art* framework, we scaled up connectomes with either homogeneous or heterogeneous interareal distributions of connectivity weights and found that only the larger reservoirs with heterogeneous wiring could overcome the lower performance inherent to the underlying connectivity. This points out once again to the importance of random wiring diagrams for ESNs’ performance. This fact is as well in line with a recent study using human connectivity as reservoir of ESNs, which showed that random connectivity indeed achieved globally maximal performances across almost all tested hyperparameters, provided the wiring cost is not considered [24].

The functional importance of randomness is also consistent with the fact that stochastic processes play a fundamental role in brain connectivity formation, both at a micro and meso/macro-scale, as supported by empirical [25], and computational modeling studies [26, 27].

While here we tested the performance of the ANNs in two memory tasks, our approach is versatile and extendable, since it allows an open ended examination of the consequences of network topology found in nature for artificial systems. Specifically, the following contributions hold: First, we offer an approach for creating ANNS with network topology dictated directly from empirical observations in BNNs. Second, creating and upscaling BioESNs from real connectomes is in itself a highly non-trivial problem and here we offer, although not exhaustively, insights into the consequences of each strategy. Third, our method allows building ANNs with network topologies based on empirical data from diverse biological species (mammalian brain networks).

We are aware of a number of limitations of our study as well as interesting research avenues for future work.

We evaluated our BioESNs models on two different memory tasks framed as regression problems; so future work could, for example, include classification tasks as well as more ecologically realistic tasks.

Connections in the adult brain change constantly as a consequence of stochastic fluctuations and activity-driven plasticity, e.g., learning and memory [28]. In our study, we assumed connectivity within the reservoir to be constant during the tasks. Previous studies have shown some effects of plasticity rules on ESNs [29], so we foresee interesting future work along those lines as well.

As we aimed at testing the potential impact of the global wiring diagram of connectomes, we consider the entire connectomes as one unique network to create the reservoirs. This is different from a previous study where the connectivity was divided into subnetworks corresponding to brain systems that were separately trained [24]. We decided to avoid here the strong assumptions that such an approach implies, but we recognize a potential for future studies in the direction, for example, exploring the division of networks as different input/output subsystems.

## Conclusion

The wiring of biological and artificial neural networks plays a crucial role in providing networks with fundamental built-in biases that influence the their ability to learn and their performance. Brain connectivity results from emergent complex phenomena involving evolution, ontogenesis and plasticity, while artificial neural networks are deliberately hand-crafted. Our presented work represents a new interface between network neuroscience and artificial neural networks, precisely at the level of connectivity. We contribute an original approach to blend real brain connectivity and artificial networks, paving the way to future hybrid research, a promising exploration path leading to potential better performance and robustness of artificial networks and understanding of brain computation.

## Materials and methods

### Echo State Networks (ESN)

Echo State Networks (ESNs) are one kind of recurrent neural networks (RNNs) belonging to the broader family of reservoir computing models, typically used to process temporal data [30]. The ESN model consists of an input, a reservoir and an output layer. The input layer feeds the input(s) signal(s) into a recurrent neural network with fixed weights, i.e., the reservoir. The function of the reservoir is to non-linearly map the input signal onto a higher dimensional space by means of the internal states of the reservoir. Formally, the input vector 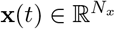 is fed into the reservoir through an input matrix 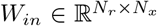, where *N_r_* and *N_x_* indicate the number of reservoir and input neurons, respectively. Optionally, the input can be scaled by a factor 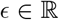 (*input scaling*) before been fed into the network. The discrete dynamics of the leaky neurons in the reservoir are represented by the state vector 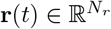 and governed by the following equations:

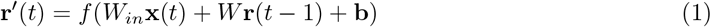

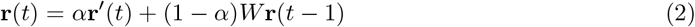

Where 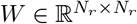 is the connectivity matrix between reservoir neurons, 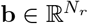 is the bias vector, *f* the nonlinear activation function.

For all the presented results *f = tanh*, the hyperbolic tangent function which bounds the values of **r** to the interval [−1, 1]. With *α* =1, there is no leakage, which we found to perform better so we fixed them for all presented results. Thus, Eq 1 and Eq 2 can be re-written together as:

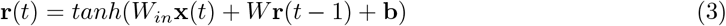

The output readout vector 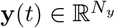 is obtained as follows:

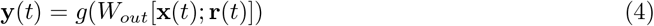

Where *g* is the output activation function and indicates the vertical vector concatenation and 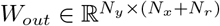 is the readout weights matrix. For all results presented *g* was either rectified linear unit (*ReLU*) or the identity function. Training the model means finding the weights of *W_out_*. Linear regression was used to solve *W_out_* = *Z^+^Y*, where *Z^+^* is the pseudoinverse of *Z* = [x(*t*); r(*t*)], i.e., the vertically concatenated inputs and reservoir states for all time steps.

We initialized the incoming weights in *W_in_* with random uniformly distributed values between [-1,1]. Further considerations about weights initialization as well as sparsity of the reservoir are detailed in the section Mappping and upscaling connectomes with *bio2art*.

The activity of the reservoir neurons is initialized with r(*t*) = 0. That produces an initial transient of spurious activity which is unrelated to the inputs and is therefore useless for learning the relationship to the outputs. We discarded that initial transient of 100 time steps in all cases, both for training and for testing.

All presented results with ESNs training where obtained using the Python package *echoes*, publicly available [31].

### ESN hyperparameters tuning

The typically most influential hyperparameters in ESNs are reservoir size *N_r_*, spectral radius of the reservoir *ρ*, input scaling factor *ε* and the leakage rate *α* [15]. In our scheme, the reservoir connectivity W is determined by the real connectome, thus determining a fixed *N_r_*. So the hyperparameters explored were: spectral radius of the reservoir connectivity matrix *ρ* = {0.91, 0.93,…, 0.99}, input scaling *ε* = {10^-9^, 10^-8^,…, 10^0^}, leakage rate *α* = {0.6, 0.8, 1} and bias *b* = {0, 1}. A train/validation/test split of the data was performed. For each hyperparameters constellation, the model was trained on the training set and based on the validation score we chose and for the sake of comparison between different conditions, we fixed a common, not necessarily optimal but generally well-performing, set of hyper-parameters: Spectral radius ρ = 0.99, Input scaling ε = 10^-5^, Leakage rate α =1, Bias b = 1. Since the output values Memory Sequence Task are bounded to be greater than 0, we used ReLU as activation out function. Given that such boundary does not exist for the outputs of the Memory Capacity Task, we simply used the Identity as activation out function.

The data for train/validation was split as follows:

Sequence Recall Task: 800 trials for training and 200 for test for each hyperparameters/task difficulty/reservoir generation constellation.

Memory Capacity Task: 4000 time steps for training and 1000 for test for each hyperparameters/task difficulty/reservoir generation constellation (see Supporting Information). For each constellation, we tested 10 independent runs with newly instantiated networks.

After fixing the best hyperparameters, newly instantiated networks were generated and evaluated on the test set not yet seen by any model. The presented results in the main text are the test performances.

### Mappping and upscaling connectomes with *bio2art*

We refer to a *connectome* as the map of all the connections obtained from a single brain [32]. We used the following publicly available datasets: Macaque monkey [22], Marmoset monkey [23] and Human [10].

For the sake of clarity, let us disect the connectivity into two components: topology and weights. The topology refers to the wiring diagram (i.e., who connects to whom), regardless of the strength of the connection (assuming non-binary connectivity). So if we think of the connectivity in terms of its representations as a connectivity matrix, the topology refers here to the binary mask that indicates which positions of the matrix have values different from zero. The weights describe the precise strength of those connections between neurons. This differentiation is not necessarily completely consistent with common uses in the literature, but serves the purpose of explaining the work presented here. As our goal is to evaluate the role of the topology of real brains, i.e., the mentioned wiring diagram, we propose a scheme to map real connectomes onto reservoirs of ESNs, with topology corresponding to real brains, but weights drawn from a uniform distribution of values between [-1, 1], as in classical ESN approaches [30]. Classical ESNs reservoir have weights randomly from a symmetric probability distribution, typically Uniform or Gaussian, and place them at random between neurons, thus generating random graph from the perspective of the topology as well. Another common practice is to use a relatively sparse network, e.g., common choices are pairwise probability of connection *p* < 0.1 or a low fixed mean degree, e.g., *k* = 10). So for the sake of comparison and testing the effect of topology in the performance of ESNs, we the following surrogate connectivity variations as null models (see Fig. 2 for a visual comparison):

- ***Bio (rank):*** Preserves the empirical topology, i.e., wiring diagram or “who connects to whom”. Weights are placed such that the rank order of them is the same as the empirical, i.e., strong weights in the empirical connectome will correspong to higher positive weights in the *Bio (rank*) condition, and viceversa.
- ***Bio (no-rank):*** Preserves the empirical topology, i.e., wiring diagram or “who connects to whom”. Weights are placed randomly, so no rank order is preserved.
- ***Random (density):*** Wiring diagram is completely random, but allowing only as many connections as to match the density of connections of the empirical connectome. The density is defined as the fraction of present connections out of all the possible ones. Weights are placed randomly.
- ***Random (k):*** Wiring diagram is completely random, but allowing only a fixed number of connections k per neuron. All presented experiments use k = 10. Weights are placed randomly.
- ***Random (full):*** Wiring diagram is completely random and all neurons connect to all other neurons, i.e., the density of connections is 1. Weights are placed randomly.

The *bio2art* functionality builds artifical recurrent neural networks by using the topology dictated by empirical neural networks and by extrapolating from the empirical data to scale up the artifical neural networks.

We explored here a range of network size scaling factors between 1 and 30x by step of 1.

*bio2art* offers the possibility to control the within and between area connectivity as well. There are currently no empirical comprehensive data for neuron-to-neuron connectivity within each brain region. However, existing empirical data suggest that within-region connectivity strength constitutes approximately 80% of the extrinsic between-region connectivity strength [22]. Therefore, the intrinsic, within-region connectivity in our work followed this rule. It should be noted that the number of connections that a neuron can form within neurons of the same region is controlled by a parameter dictating the percentage of connections that a neuron will form, out of the total number of connections that can be formed. Here we set this parameter to 1, that is, all connections between neurons within a region are formed. The exact details of the implementation can be found here [14], together with a freely available Python toolbox to apply the tools used here.

### Tasks

#### Memory Capacity (MC) Task

In this memory paradigm, a random input sequence of numbers **X**(t) is presented to the network through an input neuron. The network is supposed to independently learn delayed versions of the input, thus there are several outputs [16]. Each output *Y_τ_* predicts a delayed version of the input **X**(*t*) by *τ* time steps, i.e., *Y_τ_*(*t*) = *X*(*t — τ*). The values of the input signal **X** were randomly drawn from a uniform distribution, i.e., X(t) ~ *Uniform*(—0.5, 0.5). The networks were trained with 4000 time steps and tested on the subsequent 1000. Each output is trained independently and the performance, the so called *Memory Capacity (MC*), is calculated as the cumulative score (squared Pearson correlation coefficient *ρ*) across all outputs (i.e., all time lags) as follows:

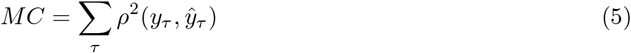

#### Sequence Recall Task

In this task the network is presented with two inputs, *X*_1_(*t*), *X*_2_(*t*), a sequence of random numbers to memorize and a cue input, respectively. The cue input signals whether to fixate (output equal to zero) or recall. After the recall signal, the network is supposed to output the memorized sequence in the L steps previous to recall signal, where the pattern length L is a parameter regulating the task difficulty (see Fig. 6). One trial of the task consists of one fixation period and the subsequent recall period. The values of the input signal *X*_1_(*t*) were randomly drawn from a uniform distribution, i.e., *X*_1_(*t*) ~ *Uniform*(0,1). The performance was evaluated with the *R*^2^ score only during the recall steps because the fixation phase was much easier for the model to get right and would have inflated the performance. Each BioESN was trained with 800 trials and tested on 200 trials. For each pattern length L in {5, 6, 7, …, 25}, 100 different networks with newly instantiated weights were tested.

## Acknowledgments

We thank Dr. Fatemeh Hadaeghi and Dr. Dong Li for fruitful discussions about ESNs and constructive feedback on the manuscript. FD has been funded by the Deutscher Akademischer Austausch Dienst (DAAD). AG has been funded by the Deutsche Forschungsgemeinschaft (DFG) (HI 1286/7-1). CCH has been funded by the Deutsche Forschungsgemeinschaft (DFG) (SFB 936/A1, Z3; TRR 169/A2; SPP 2041, HI 1286/6-1) and the Human Brain Project (SGA2, SGA3).

