## Supplemental information for "Brain Connectivity meets Reservoir Computing"

### Supporting information

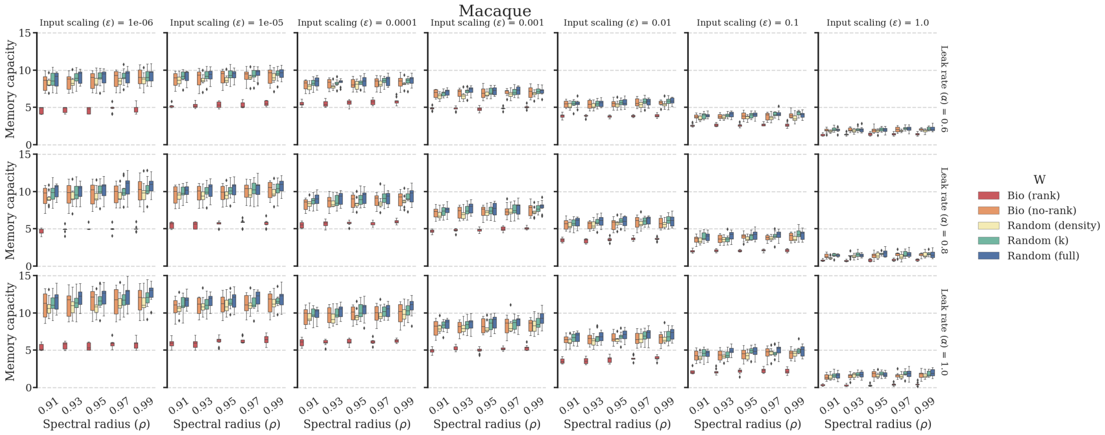


**Fig 7. Memory Capacity task. Echo state network hyperparameters grid search. Homogeneous interareal connectivity. Macaque connectome.** Results of grid search over input scaling, leakage rate and spectral radius hyperparameters. For each parameter constellation, the boxplots show the aggregated validation scores of ten independently generated and trained reservoirs.

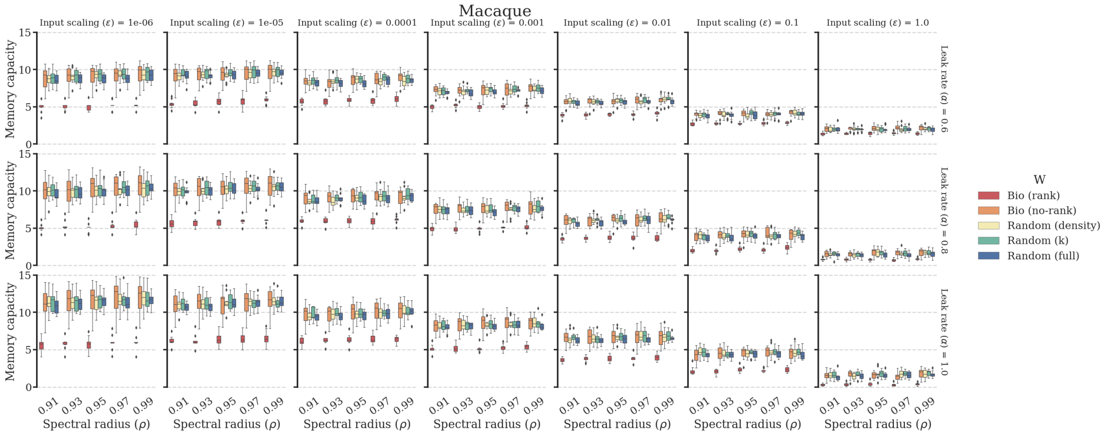


**Fig 8. Memory Capacity task. Echo state network hyperparameters grid search. Heterogeneous interareal connectivity. Macaque connectome.** Results of grid search over input scaling, leakage rate and spectral radius hyperparameters. For each parameter constellation, the boxplots show the aggregated validation scores of ten independently generated and trained reservoirs.

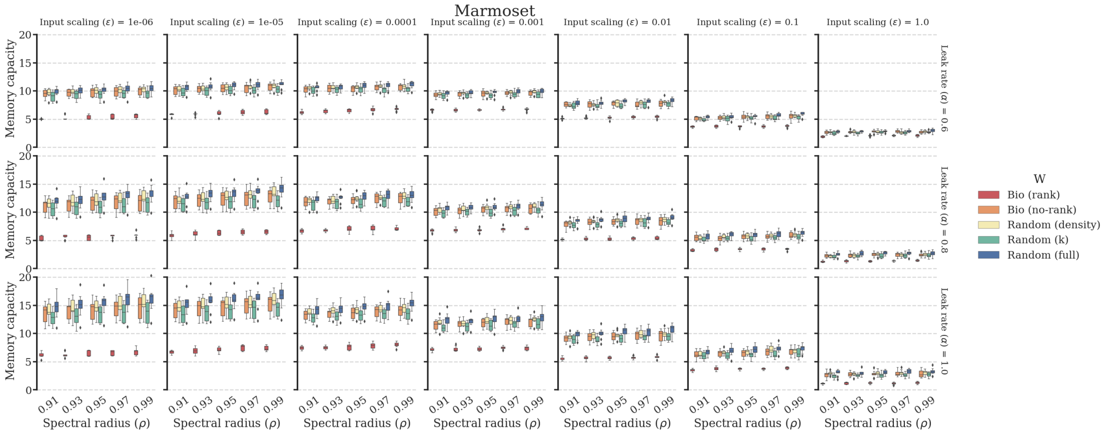


**Fig 9. Memory Capacity task. Echo state network hyperparameters grid search. Homogeneous interareal connectivity. Marmoset connectome.** Results of grid search over input scaling, leakage rate and spectral radius hyperparameters. For each parameter constellation, the boxplots show the aggregated validation scores of ten independently generated and trained reservoirs.

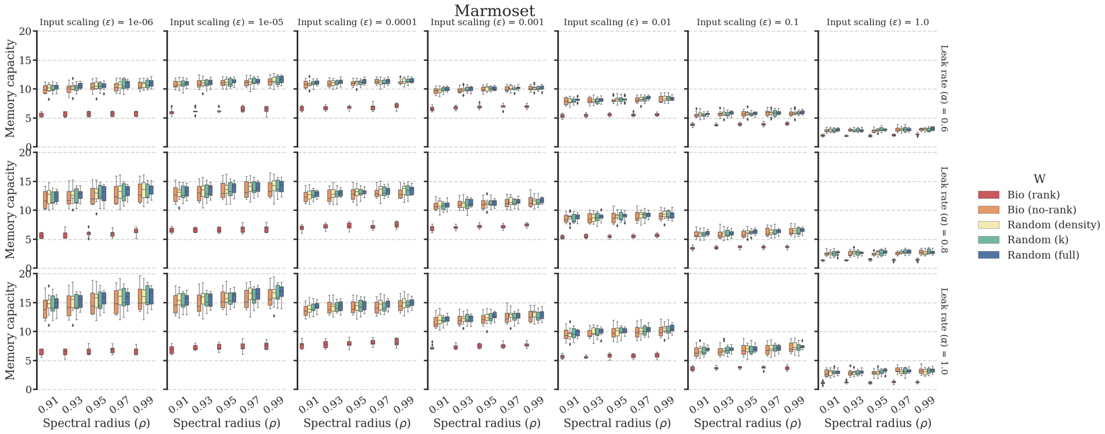


**Fig 10. Memory Capacity task. Echo state network hyperparameters grid search. Heterogeneous interareal connectivity. Marmoset connectome.** Results of grid search over input scaling, leakage rate and spectral radius hyperparameters. For each parameter constellation, the boxplots show the aggregated validation scores of ten independently generated and trained reservoirs.

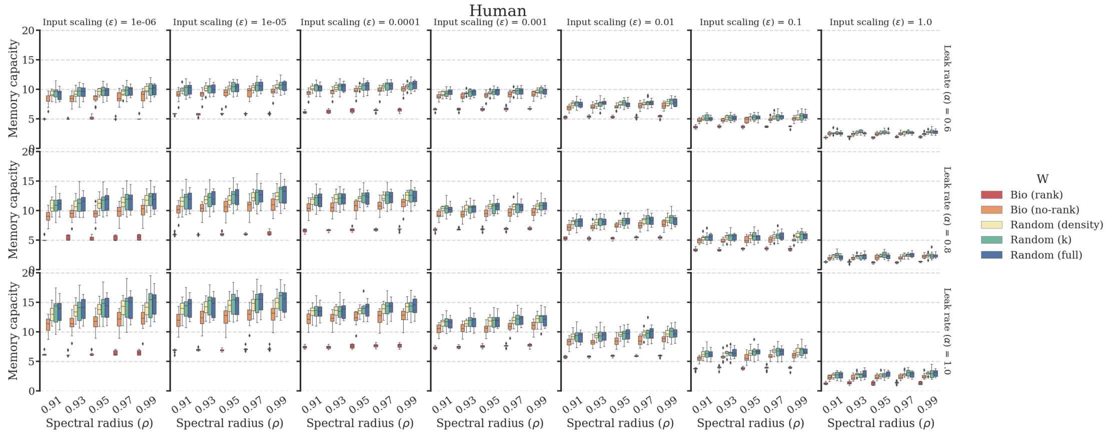


**Fig 11. Memory Capacity task. Echo state network hyperparameters grid search. Homogeneous interareal connectivity. Human connectome.** Results of grid search over input scaling, leakage rate and spectral radius hyperparameters. For each parameter constellation, the boxplots show the aggregated validation scores of ten independently generated and trained reservoirs.

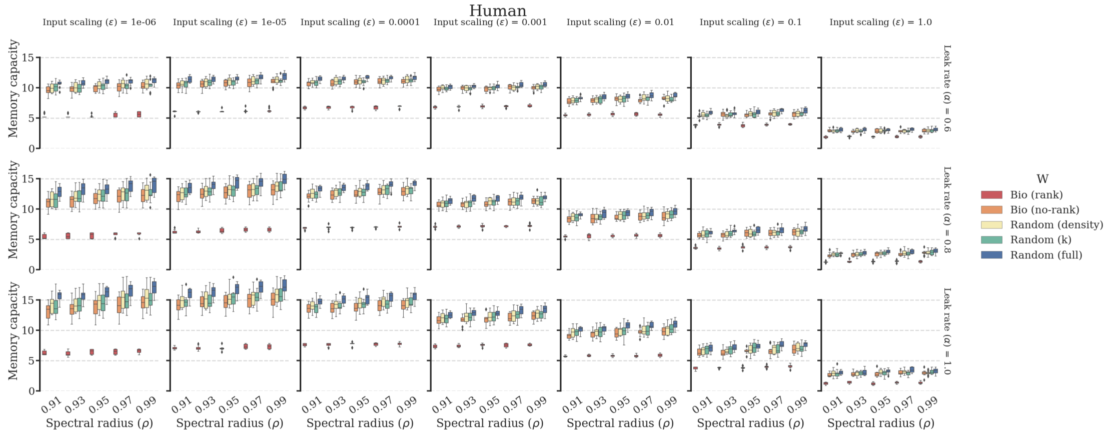


**Fig 12. Memory Capacity task. Echo state network hyperparameters grid search. Heterogeneous interareal connectivity. Human connectome.** Results of grid search over input scaling, leakage rate and spectral radius hyperparameters. For each parameter constellation, the boxplots show the aggregated validation scores of ten independently generated and trained reservoirs.
